# Attenuated beta rebound to proprioceptive afferent feedback in Parkinson’s disease

**DOI:** 10.1101/334565

**Authors:** Mikkel C. Vinding, Panagiota Tsitsi, Harri Piitulainen, Josefine Waldthaler, Veikko Jousmäki, Martin Ingvar, Per Svenningsson, Daniel Lundqvist

**Affiliations:** NatMEG, Department of Clinical Neuroscience, Karolinska Institutet, Stockholm, Sweden.; Section of Neurology, Department of Clinical Neuroscience, Karolinska Institutet, Stockholm, Sweden; Aalto NeuroImaging, Department of Neuroscience and Biomedical Engineering, Aalto University School of Science, Espoo, Finland; Cognitive Neuroimaging Centre, Nanyang Technological University, Singapore

## Abstract

Motor symptoms are defining traits in the diagnosis of Parkinson’s disease (PD). A crucial component in motor function and control of movements is the integration of efferent signals from the motor network to the peripheral motor system, and afferent proprioceptive sensory feedback. Previous studies have indicated abnormal movement-related cortical oscillatory activity in PD, but the role of the proprioceptive afference on abnormal oscillatory activity in PD has not been elucidated. In the present study, we examine the role of proprioception by studying the cortical processing of proprioceptive stimulation in PD patients, ON/OFF levodopa medication, as compared to that of healthy controls (HC). We used a proprioceptive stimulator that generated precisely controlled passive movements of the index finger and measured the induced cortical oscillatory responses following the proprioceptive stimulation using magnetoencephalography (MEG). Both PD patients and HC showed a typical initial mu/beta-band (8–30 Hz) desynchronization during the passive movement. However, the subsequent beta rebound after the passive movement that was apparent in HC was much attenuated and almost absent in PD patients. Furthermore, we found no difference in the degree of beta rebound attenuation between patients ON and OFF levodopa medication. Our results hence demonstrate a disease-related deterioration in cortical processing of proprioceptive afference in PD, and further suggest that such disease-related loss of proprioceptive function is due to processes outside the dopaminergic system affected by levodopa medication.

## 1 Introduction

Parkinson’s disease (PD) is a common progressive neurodegenerative disease. The diagnosis of PD is based on the presence of bradykinesia, tremor, and rigidity. The motor symptoms are mainly caused by deficient dopamine neurotransmission and are counteracted by dopamine replacement therapies. (Kalia & Lang, 2015). The pathology of PD is characterized by the presence of alpha-synuclein-enriched protein aggregates called Lewy-bodies (Kalia & Lang, 2015; Rodriguez-Oroz et al., 2009). Lewy body pathology spreads progressively from the olfactory bulb and brainstem to larger parts of the brain giving rise to motor symptoms but also non-motor symptoms such as hyposmia, depression, sleep disorders, pain, and cognitive impairment (Braak et al., 2003; Kalia & Lang, 2015).

One of the core symptoms of PD is disturbances in proprioception crucial for successful control of movements and for maintaining balance and posture (Dietz, 2002; Konczak et al., 2009). Proprioceptive signals are afferent neural signals from the peripheral nervous system (PNS) to the central nervous system (CNS). The proprioceptive signals primarily originate in the muscle spindles, Golgi tendon organ, and joint receptors, and project through the spinal cord to the dorsal spinocerebellar tract and from there to the cerebellum, thalamus, and further to the sensory-motor areas in the cortex (Proske & Gandevia, 2012). Carrying out voluntary movements requires efferent signals from cortex through basal ganglia through the brainstem to the PNS, and afferent proprioceptive signals going back, transmitting information about the prior state of the locomotor system and its changes, e.g., in limb position.

Disturbances in proprioception in PD is a central factor in the development of the motor symptoms in PD (Dietz, 2002; Konczak et al., 2009). PD patients are, for instance, worse compared to healthy controls (HC) at detecting passive movements of their limbs which is dependent on proprioceptive afferents (Konczak, Krawczewski, Tuite, & Maschke, 2007; Maschke, Gomez, Tuite, & Konczak, 2003; Zia, Cody, & O’Boyle, 2000). The apparent deterioration in the utilization of proprioceptive information in PD does not appear to be caused by disturbances in the PNS. Recordings of muscle spindle responses by microneurography show no differences in afferent signals between HC and PD patients (Mano, Yamazaki, & Mitarai, 1979). The early cortical processing of proprioceptive signals— measured with electroencephalography (EEG) as the event-related potentials (ERPs) following passive movements of the index fingers—does not differ between PD patients and HC (Seiss, Praamstra, Hesse, & Rickards, 2003).

Behavioral studies indicate that loss of proprioceptive function arises due to errors in central processing and sensory integration. For example, illusions of movement induced by vibrating muscles in the limbs are reduced in PD patients compared to HC (Rabin, Muratori, Svokos, & Gordon, 2010; Rickards & Cody, 1997). PD patients also show increased dependency on visual cues over proprioceptive feedback during active movements (Demirci, Grill, McShane, & Hallett, 1997; Nowak & Hermsdörfer, 2006), and postural control is more difficult for PD patients without visual feedback compared to HC (Adamovich, Berkinblit, Hening, Sage, & Poizner, 2001; Jacobs & Horak, 2006).

The disturbances in proprioception in PD thus appear to arise in the higher levels of sensorimotor integration. Impaired utilization of proprioceptive information in PD patients also shows when switching from visually guided to the proprioceptive guided control of balance (Bronstein, Hood, Gresty, & Panagi, 1990). Errors in integrating proprioceptive signals are seen in grasping tasks where PD patients show increased grip force when grasping objects compared to HC, suggesting that proprioceptive feedback and active motor commands are not adequately integrated to facilitate optimal grasping (Fellows, Noth, & Schwarz, 1998; Nowak & Hermsdörfer, 2006). Impaired proprioception degrades sensorimotor integration, which is compensated with feedback from other sensory domains (Abbruzzese & Berardelli, 2003; Konczak et al., 2009).

Disturbances in the proprioceptive processing in PD appears to be due to errors in the integration of proprioceptive afferents, but the actual mechanisms of the disturbances in the processing of proprioceptive signals are unknown. If loss of proprioception in PD is due to disturbed communication between the basal ganglia, thalamus, and cortical motor areas, due to a faulty integration of proprioceptive signals later at a later processing stage, or outside the dopamine-dependent pathways has not yet been adequately explained (Rabin et al., 2010). Isolating the relative contribution from efferent and afferent signals on movement control to answer how afferents are affected in PD is difficult, as both are necessary for successfully carrying out the movements and depends on functional and anatomical overlapping neural processes (Prud’homme & Kalaska, 1994). In the present study, we investigate how the contribution from afferent proprioceptive afferents are processed in PD in the absence of efferent motor signals by stimulating only the proprioceptive afferents in passive movements.

PD is associated with changes in neural oscillatory behavior demonstrated both at local and global levels. Local field potentials from the subthalamic nucleus (STN) in PD patients show an increase in beta-band (~14–30 Hz) oscillations (Alonso-Frech et al., 2006). The abnormal beta oscillations are decreased by administration of dopaminergic medication, indicating a functional link between dopamine levels and background beta oscillations (Alonso-Frech et al., 2006; Neumann et al., 2017). The decrease of beta-band oscillations in STN due dopaminergic medication have been correlated to an overall reduction in motor symptoms in PD (Kühn, Kupsch, Schneider, & Brown, 2006). Although PD is associated with an increase of beta-oscillations in sub-cortical areas, cortical beta oscillations are decreased in PD patients compared to HC (Bosboom et al., 2006; Heinrichs-Graham, Kurz, et al., 2014). While cortical beta-oscillations are decreased in PD, several studies report an increase in the functional connectivity in the beta-band within the sensory-motor cortex (Bosboom, Stoffers, Wolters, Stam, & Berendse, 2009; Heinrichs-Graham, Kurz, et al., 2014; Pollok et al., 2013; Silberstein et al., 2005). The level of synchronicity of cortical beta activity is related to rigidity and action tremor in PD (Airaksinen et al., 2012).

Cortical beta-band oscillations are actively involved in sensorimotor processing. Beta-band oscillations exhibit well-known event-related desynchronization (ERD) and event-related synchronization (ERS) during active states of the sensorimotor system (Fig. 1). Beta oscillations attenuate in the second before movement onset, known as the movement-related ERD, and is prevalent during the duration of the movement (Cheyne, 2013; Jurkiewicz, Gaetz, Bostan, & Cheyne, 2006; Kilavik, Zaepffel, Brovelli, MacKay, & Riehle, 2013; Salmelin & Hari, 1994). Once the movement stops, the beta oscillations temporarily show a relative increase, known as the post-movement ERS or *beta rebound*, before going settling back at the baseline level. (Cheyne, 2013; Kilavik et al., 2013; Neuper & Pfurtscheller, 1996; Pfurtscheller, Stancák, & Neuper, 1996; Salmelin & Hari, 1994). The origin of both the movement-related beta ERD and the beta rebound during voluntary movements is the primary somatosensory cortex (Druschky et al., 2003; Xiang et al., 1997) with the cortical source of the movement-related being more posterior than the source of subsequent beta rebound (Jurkiewicz et al., 2006).

**Figure 1:**
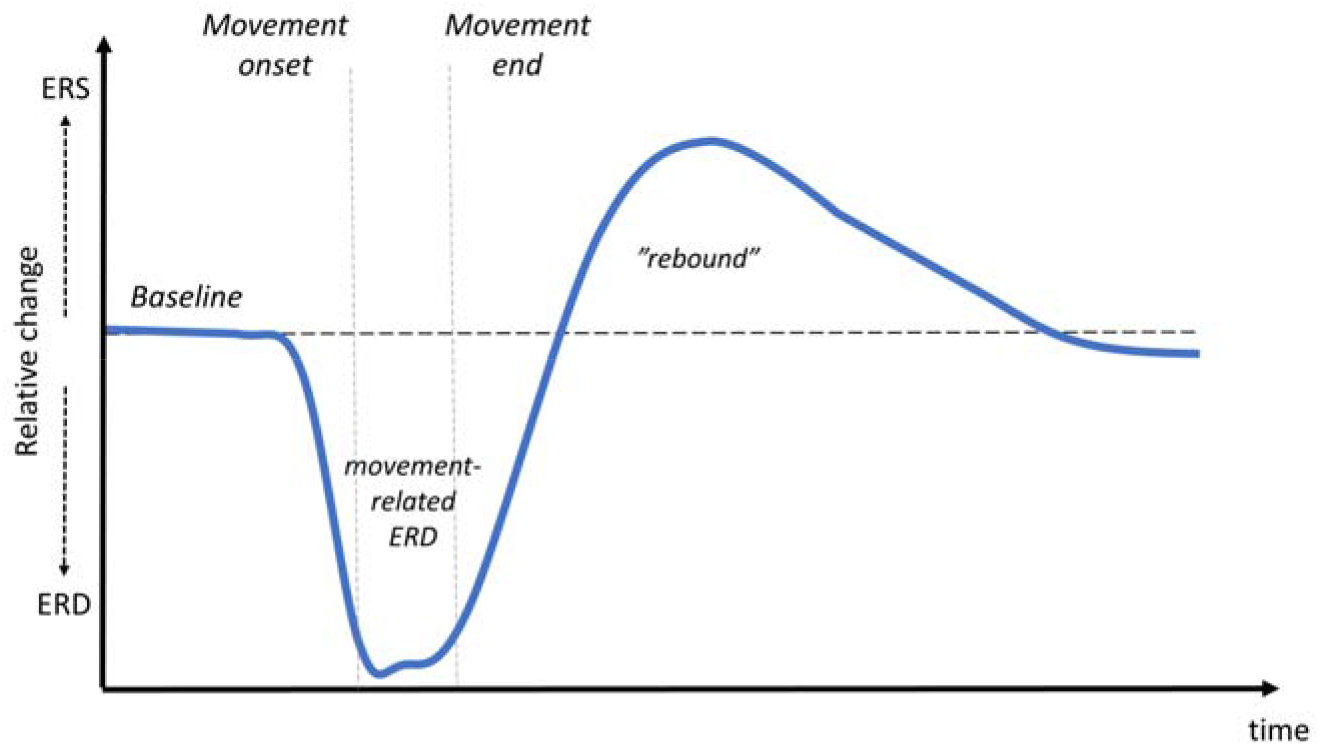
Movement-related beta-band activity. Typical event-related synchronization (ERS) and desynchronization (ERD) in the beta-band during movements measured from the cortex with EEG/MEG. When initiating a movement beta activity start to desynchronize and prevails as a persistent ERD during the movement execution phase. Once the movement ends, it is followed by an ERS referred to as the beta rebound.

The movement-related ERD and beta rebound seen during voluntary movements is attenuated in PD compared to HC. PD patients show less beta ERD before and during active movements and a smaller rebound after active movements (Devos et al., 2003; Heinrichs-Graham, Wilson, et al., 2014; Pfurtscheller, Pichler-Zalaudek, Ortmayr, Diez, & Reisecker, 1998). It is currently unclear if the attenuated dynamics in the beta-band in PD is driven by deficits in efferent processes, processes of afferents, or at a higher level in the integration of afferent and efferent signals, as both efferent and afferent signals, and the integration of the two, is needed for carrying out voluntary movements. The movement-related ERD is taken to reflect the active state of the motor system, receiving the afferent signals, and the cortical processes responsible for integrating the efferent and afferent signals (Engel & Fries, 2010).

The beta rebound has been linked to proprioceptive afferents. The beta rebound is present for passive induced movements in healthy subjects (Alegre et al., 2002; Parkkonen, Laaksonen, Piitulainen, Parkkonen, & Forss, 2015) meaning the beta rebound is related to the processing of proprioceptive signals independent of the efferent commands from cortex to the periphery. The beta rebound is also diminished when applying a temporary ischemic nerve block to disrupt the proprioceptive feedback from the muscles in healthy subjects (Cassim et al., 2001). We hypothesize that the attenuated beta ERD and rebound during voluntary movement in PD to some extent related to disturbances in the cortical processing of proprioceptive signals—even though the relative contribution of efferent signals, afferent signals, and the integration is unclear.

In the present study, we investigate the differences between PD patients compared to HC regarding the cortical processing of proprioceptive information. In contrast to earlier studies that have used voluntary movements to study beta-oscillations in PD, we isolate the processing of *afferent* from that of *efferent* information by using a computer-controlled proprioceptive stimulator that generates precise passive movements of the index finger. With this method, we examine the processing of proprioceptive signals in isolation, without the confounding effect of efferent motor signals. If the attenuation of movement-related ERD and rebound during active movements is due to defect in the processing of proprioceptive afferents in PD, we expect the movement-related ERD and beta rebound to be less salient for PD patients than for HC. Conversely, if the difference between PD and HC primarily depends upon deficits in *efferent* signaling processing, then we expect there to be no differences between PD and HC during passive movements. We furthermore investigated how Levodopa influences processing of proprioceptive information in PD patients, by examining PD patients both in ON and OFF Levodopa states. While Levodopa improves motor symptoms in PD, it is unclear to what extent it affects proprioceptive processing, as results from behavioral studies have been inconclusive with regards to improvement of proprioceptive function due to medication (Konczak et al., 2009).

## 2 Materials and methods

### 2.1 Participants

Thirteen PD patients (age 41–75; three female) and seventeen HC (age 54–76; five female) participated in the study. One patient had to abort the session due to severe tremor in the OFF-medication state and subsequently had to cancel the participation in the study. One HC was excluded from analysis due to insufficient quality of the MEG recording.

The inclusion criteria for the PD group were: a clinical diagnosis of PD according to the United Kingdom Parkinson’s Disease Society Brain Bank Diagnostic Criteria with Hoehn and Yahr stage 1 to 3 (Hoehn & Yahr, 1967), and treatment with Levodopa, COMT inhibitors, MAO-B inhibitors, or dopamine receptor agonists. Additional inclusion criteria were: age between 40 to 80 years, normal according to a physical examination (excluding parkinsonism), adequate cognitive status in terms of well-functioning and being able to give written consent after being informed about the procedure and purpose of the study. Exclusion criteria were: diagnosis of major depression dementia, history or presence of schizophrenia, bipolar disorder, epilepsy, or history of alcoholism or drug addiction according to Diagnostic and Statistical Manual of Mental Disorders, fifth edition (American Psychiatric Association, 2013).

HC were recruited from a pool of healthy participants who previously had participated in other studies within the preceding year, or amongst eligible spouses of PD patients. Exclusion criteria for HC were: Diagnosis of PD or any other movement disorder, depression, dementia, the presence of- or history of schizophrenia, bipolar disorder, epilepsy, or history of alcoholism or drug addiction.

**Table 1:** Summary of the PD patient group and the HC group

| | PD patients | Healthy controls | BF ($H_1/H_0$) |
| --- | --- | --- | --- |
| N | 12 | 16 |  |
| Sex | 3 female, 9 male | 5 female, 11 male | 0.42 |
| Age | 44-75 years (mean: 63.7 years) | 54-76 years (mean: 69.6 years) | 2.02 |
| Disease duration | 1-11 years (mean: 5.5 years) | - |  |
| LEDD | 632 mg (SD: 271 mg) | - |  |
| MDS-UPDRS-III OFF | 31.0 (SD: 13.2) | 1.1 (SD: 1.7) | $5.36 \cdot 10^6$ |
| MDS-UPDRS-III ON | 16.3 (SD: 10.2) | - |  |
| MoCA | 25.7 (SD: 3.1) | 26.1 (SD: 1.9) | 0.30 |
| HADS Anxiety | 4.1 (SD: 3.1) | 2.9 (SD: 1.7) | 0.52 |
| HADS Depression | 2.9 (SD: 2.6) | 1.6 (SD: 1.0) | 1.23 |

The local ethics committee in Stockholm approved the study, and we obtained signed consent from participants following the Declaration of Helsinki. The patients were recruited from the Parkinson’s Outpatient Clinic, Department of Neurology, Karolinska Huddinge University Hospital (Stockholm, Sweden).

### 2.2 Proprioceptive stimulation

The proprioceptive stimulation consisted of passive movements of the index finger on the hand contralateral to PD dominant side for PD patients and on the dominant hand for HC.

The passive finger movements were evoked by a custom-made MEG compatible pneumatic movement actuator utilizing a pneumatic artificial muscle (PAM) (Piitulainen, Bourguignon, Hari, & Jousmäki, 2015). The PAMs contract when filled with pressurized air and expand when air-pressure is released, resulting in movement along a single axis. The flow of pressurized air to the device was controlled from outside the magnetically shielded room using Presentation software (v. 18.3; www.neurobs.com). The induced movements consisted of one contraction and extension of the pneumatic artificial muscle with a 200 ms interval (see Fig. 2B) inducing a passive finger movement with an amplitude of 0.5 cm. Each movement was followed by a silent period between 3.5 s to 4.0 s until the next induced movement. Each session contained a total of 90 induced movements.

**Figure 2:**
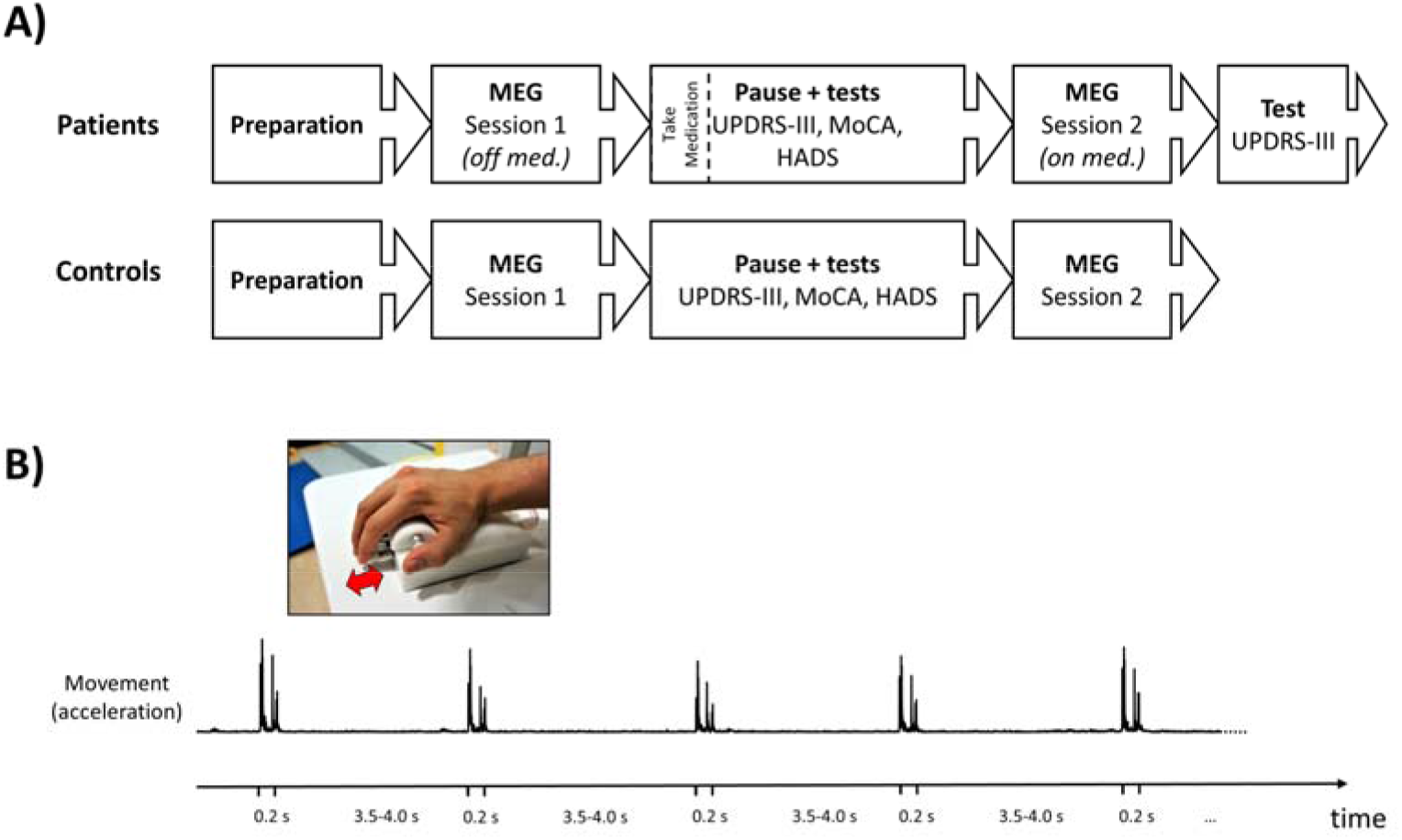
Experimental procedure and task. A) Overview of the experimental procedure. B) In the task, subjects had their index finger on a pneumatic artificial muscle (inserted picture) that contracted and substrated within a 200 ms interval inducing passive movements once every 3.5-4 seconds—illustrated by the continuous acceleration measured by an accelerometer attached to the index finger.

### 2.3 Procedure

We tested each PD patient in OFF (at least 12 hours after last levodopa intake) and ON (one hour after levodopa intake) medication status in two consecutive sessions on the same day. After collecting consent from participants, subjects were then prepared for MEG and placed in the MEG scanner inside a magnetically shielded room (MSR). While seated in the scanner, subjects received proprioceptive stimulation by induced passive movements as described above. The subjects were watching a movie with sound throughout the proprioceptive stimulation.

MEG recordings in the MSR were acquired twice: first in OFF medication state and then after an hour’s break in ON. The motor subscale of the Movement Disorder Society’s Unified Parkinson’s Disease Rating Scale (MDS-UPDRS part III; Goetz et al., 2007) was assessed in both ON and OFF, while the Montreal Cognitive Assessment battery (MoCA) and the Hospital Anxiety and Depression Scale (HADS) were assessed in ON. A neurologists certified in the use of MSD-UPDRS did all assessments.

HC was tested twice to accommodate the effect of time between repeated sessions (Wilson, Heinrichs-Graham, & Becker, 2014). The experiment was repeated in HC with the same hour distance break as PD subjects but did not include any PD medication for HC. By repeating the assessment on both PD patients and HC, we were able to isolate the effect of the medication this being the only difference between the two recordings among our groups.

### 2.4 MEG data acquisition

MEG data were recorded with an Elekta Neuromag TRIUX 306-channel MEG system, with 102 magnetometers and 204 planar gradiometers. The MEG scanner was located inside a two-layer magnetically shielded room (model Ak3B from Vacuumschmelze GmbH). Data were sampled at 1000 Hz with an online 0.1 Hz high-pass filter and 330 Hz low-pass filter. Internal active shielding was active to suppress electromagnetic artifacts from the surrounding environment. Subjects head position and head movement inside the MEG helmet were sampled with MEG data using head-position indicator coils (HPI). The HPI was attached to subjects’ heads, and the location of the HPI location was digitalized with a Polhemus Fastrak motion tracker.

Horizontal and vertical electrooculogram (EOG) and electrocardiogram (ECG) were recorded simultaneously with bipolar Ag/AgC electrodes located above/below the left eye (vertical EOG) and on each side of the eyes (horizontal EOG). Electromyography (EMG) was measured on the forearms above the flexor carpi radialis with bipolar Ag/AgC electrodes located 7-8 cm apart. The position of the EMG electrodes was determined by asking subjects to tap their fingers and then locate muscle movements. An accelerometer (ADXL335; Analog Devices Inc., Norwood, MA) attached to the nail of the index finger measured the acceleration of the finger movements along three orthogonal axes. Continous time-course of the accelerometer were sampled together with the MEG data.

### 2.5 Data processing

#### 2.5.1 Pre-processing

MEG data were processed off-line first by applying temporal signal space separation (tSSS) to suppress artifacts from outside the scanner helmet and correct for head movement during the recordings (Taulu, Kajola, & Simola, 2004; Taulu & Simola, 2006). The tSSS had a buffer length of 10 s and a cut-off correlation coefficient of 0.98. Head origin was shifted to a position based on the average initial position from the first and second scan for each subject.

The continuous data from both sessions were concatenated for each subject, and independent component analysis (ICA) was then performed on the combined data using the *fastica* algorithm (Hyvarinen, 1999) implemented in MNE-Python (Gramfort et al., 2013). Components related to eye-blinks and heartbeats were identified by selecting components correlating with peaks of the measured EOG and ECG and removed from the raw MEG data. The ICA cleaned continuous MEG data was chopped into epochs from 1.5 s before movement onset to 3.5 s after movement. We rejected trials with extreme jump-artifacts based on min-to-max peak range exceeding 10 pT for the magnetometer and exceeded 2000 fT/cm for gradiometers.

The accelerometer data was filtered with a band-pass filter between 1–195 Hz, before averaging the three orthogonal channels by calculating the Euclidian norm. Trials in which subjects made accidental movements were rejected from the analysis. Accidental movements were defined as movement measured by the accelerometer outside the time window from 0–0.5 s relative to stimulus onset.

The MEG data in the remaining epochs (range 66–90, median=87) were averaged per session for each subject. The averaged response was subtracted from every single trial for each subject, to enhance sensitivity to non-phase locked responses (Kalcher & Pfurtscheller, 1995; Pfurtscheller & Lopes da Silva, 1999).

The EMG data from the forearms were cut into epochs corresponding to the time-windows of interest for the MEG data (after applying a discrete Fourier transform filter to suppress 50 Hz line noise), and the signals were rectified. The rectified EMG epochs were averaged for each session per subject. For further comparison, we calculated the power spectral density of the non-rectified EMG signals by fast Fourier transform after applying a Hann window.

#### 2.5.2 Time-frequency analysis of MEG

For the time-frequency analysis and the subsequent statistical analysis, we used the FieldTrip toolbox (Oostenveld, Fries, Maris, & Schoffelen, 2011) in Matlab R2016a (MathWorks Inc., Natick, MA). We obtained the event-related induced responses in the beta, alpha/mu, and theta bands by time-frequency decomposition using wavelets with a width of five cycles in the time window starting 1.25 s before movement onset and ending 2.5 seconds after on all frequencies from 2–40 Hz in steps of one Hz.

After time-frequency decomposition, we combined the orthogonal gradiometer pairs to get a single time-frequency representation for each gradiometer-pair location.

### 2.6 Statistical analysis

#### 2.6.1 Peripheral muscle activation

We compared the EMG activation between the different groups and sessions to ensure that subjects did not voluntarily or unknowingly move their fingers during the proprioceptive stimulation. The absence of peripheral muscle activation was taken to indicate the absence of efferent signals. To quantify whether there was any difference in peripheral muscle activation, we tested for differences in the measured EMG signals in the time-window of the time-frequency analysis.

Cluster-based permutation tests (Maris & Oostenveld, 2007) were done between each session within- and between groups on the entire time-winding of the average rectified EMG data. The tests were done by comparing each time point of the EMG time-series with a two-tailed t-test (df=26), and by summing the t-value of neighboring time-points with a p-value<0.05 (two-tailed). The sum of the clusters of t-values was compared to a distribution cluster values calculated from random permutations of data assigned to the groups using Monte Carlo simulation (n=1000). Clusters in which the total sum of t-values was on the edges of the permutation distribution beyond the critical alpha (alpha=0.05, two-tailed) were considered a significant difference.

#### 2.6.2 Beta/mu band response

The primary purpose of the study was to compare the induced responses to proprioceptive stimulation in the beta-band between PD patients and HC, and between PD patients ON/OFF Levodopa medication.

To constrain the analysis and accommodate individual differences in the position of the head inside the MEG helmet that otherwise would have reduced statistical sensitivity, we focused the statistical comparison on the combined gradiometer-pair that showed the highest amplitude in the time interval from 50 ms to 110 ms after stimulation onset. We averaged all trials across conditions per subject, after applying a 90 Hz low-pass filter and combine each orthogonal gradiometer pairs by taking the root of the squared values. The combined gradiometer-pair that showed the highest mean value across the specified time-interval in the phase-locked domain was taken to represent the sensory-motor response to the optimal proprioceptive stimulation. The peak channel was used for the statistical analysis of the induced time-frequency beta-band responses to proprioceptive stimulation.

We extracted the time-frequency response from the selected channel in the frequencies within the “mu” spectrum from 8 to 30 Hz, which encompasses the beta (14–30 Hz) and alpha (8–3 Hz) sensory motor rhythms. Comparisons of beta and mu response were combined in a single analysis due to the harmonic component of the mu rhythm, which leaks into the beta-band range (Hari, 2006).

The first effect we tested was for a general change in the beta-band response in PD compared to HC. We compared the OFF-state for PD patients to the first session for HC to get between-group comparison without additional variation introduced by medication.

The time-frequency representation of the data was log transformed and the average log-transformed power in a baseline-window from 1.25 s to 0.2 s before stimulation onset was subtracted per frequency bin from the log-transformed time-frequency data. The baseline corrected log-transformed time-frequency representation was compared between groups using a cluster-based permutations test (Maris & Oostenveld, 2007). This method for inferential statistics identifies clusters of adjacent time-frequency points along either the time or frequency domains that differ in a point-wise t-test comparison. The t-values of all points in each cluster were summed, and the sum was compared to a distribution of summed cluster values drawn from the same test in which data had been randomly sampled assigned groups using Monte Carlo simulation (n=1000). Clusters which total sum greater than 95% of the sum permutated clusters are considered a significant difference between groups.

The second effect we investigated was the effect of Levodopa medication on the beta-band response to the proprioceptive stimulation. To accommodate the effect of repetition upon the effect of medication, we tested the effect of medication in PD patients with a pseudo two-by-two design. The test-retest effect would be present for both the patient group and the HC group, but any medication specific effect would only be present in the patient group since they were the only group who did take any medication. The interaction effect between group and session would regress out the retest effect and represent the effect of the medication.

The comparison was made by subtracting the log-transformed time-frequency responses from the first and second session for each subject. The time-frequency difference was then baseline corrected by subtracting the average of the difference in a baseline from 1.25 s to 0.2 s before stimulation onset. Finally, we tested for differences between the groups on the Time-Frequency representation (TFR) differences with a cluster-based permutation test with 1000 random permutations, where clusters beyond the critical alpha (alpha=0.05, two-tailed) of the permutation distribution were considered significant.

#### 2.6.3 Baseline beta oscillations

The amount of movement-related beta-band ERS and ERD in the beta-band may depend on the spectral power in the beta-band within the baseline period leading up to the movements (Heinrichs-Graham & Wilson, 2016; Shin, Law, Tsutsui, Moore, & Jones, 2017). We tested for a relationship between the absolute power spectral density in the time-window from 1.25 s to 0.2 s before the passive movements were initiated and the relative change within clusters that showed significant differences in the primary analysis.

We tested for a relationship between baseline power and relative power change due to proprioceptive stimulation by fitting a linear regression model that explained the mean spectra power change within the cluster as a function of baseline power and regressors indicating the session, group, and individual intercepts per subject. The model was compared by Bayesian model comparison (Rouder, Morey, Speckman, & Province, 2012) to a model containing only regressors indicating the session, group, and individual intercepts, but not baseline power.

## 3 Results

### 3.1 Subject variables

Table 1 shows a summary of demographic variables (age, sex), clinical scores (disease duration, Levodopa equivalent daily dose (LEDD) (Tomlinson et al., 2010), MDS-UPDRS-III scores in ON and OFF medication status, MoCA, HADS depression and anxiety subscales), and the comparison between the two groups. PD patients and HC did not differ in the male/female ratio (BF=0.42), nor in cognitive ability measured by MoCA (BF=0.30), and anxiety score on HADS (BF=0.52). There was a trend in favor of a difference on HADS depression score (BF=1.23), with PD patients scoring higher than HC, and that HC on average was older than PD patients (BF=2.02). None of these trends are sufficient to conclude clear differences between PD patients and HC (Wetzels et al., 2011). PD patients showed an improvement of motor symptoms after taking medication reflected by the difference in MDS-UPDRS-III score between ON and OFF states (BF=4.30*10^4^).

The number of trials used in the analysis of cortical responses to the proprioceptive stimulation after data cleaning ranged between 66-90 trials with a median of 87 trials. Comparison of the number of useful trials after data cleaning in a “Bayesian ANOVA” (Rouder et al., 2012) showed there were no differences between session (BF_H1/H0_=0.33), between groups (BF_H1/H0_=0.45) or in the interactions between session and group (BF_H1/H0_=0.062).

### 3.2 Peripheral muscle activation

None of the permutation tests on the EMG time-series showed significant differences between groups (first session: p=0.80, second session: p=0.84) or between sessions (p=0.12 for PD patients and p=0.68 for HC). None of the subjects showed time-locked muscle activation to the passive finger movements, confirming that no efferent signals occurred.

### 3.3 Time-frequency responses in beta/mu band

The time-frequency responses to the proprioceptive stimulation in the beta and mu band showed a significant difference between PD patients OFF medication and HC. The significant difference was defined by a single cluster located 1.0 s after stimulation onset, lasting 0.5 s, covering a frequency range from 14 Hz to 25 Hz (p=0.017). The cluster corresponds to the post-movement beta rebound, which was considerably attenuated, almost absent, in PD patients compared to HC (Fig. 3).

**Figure 3:**
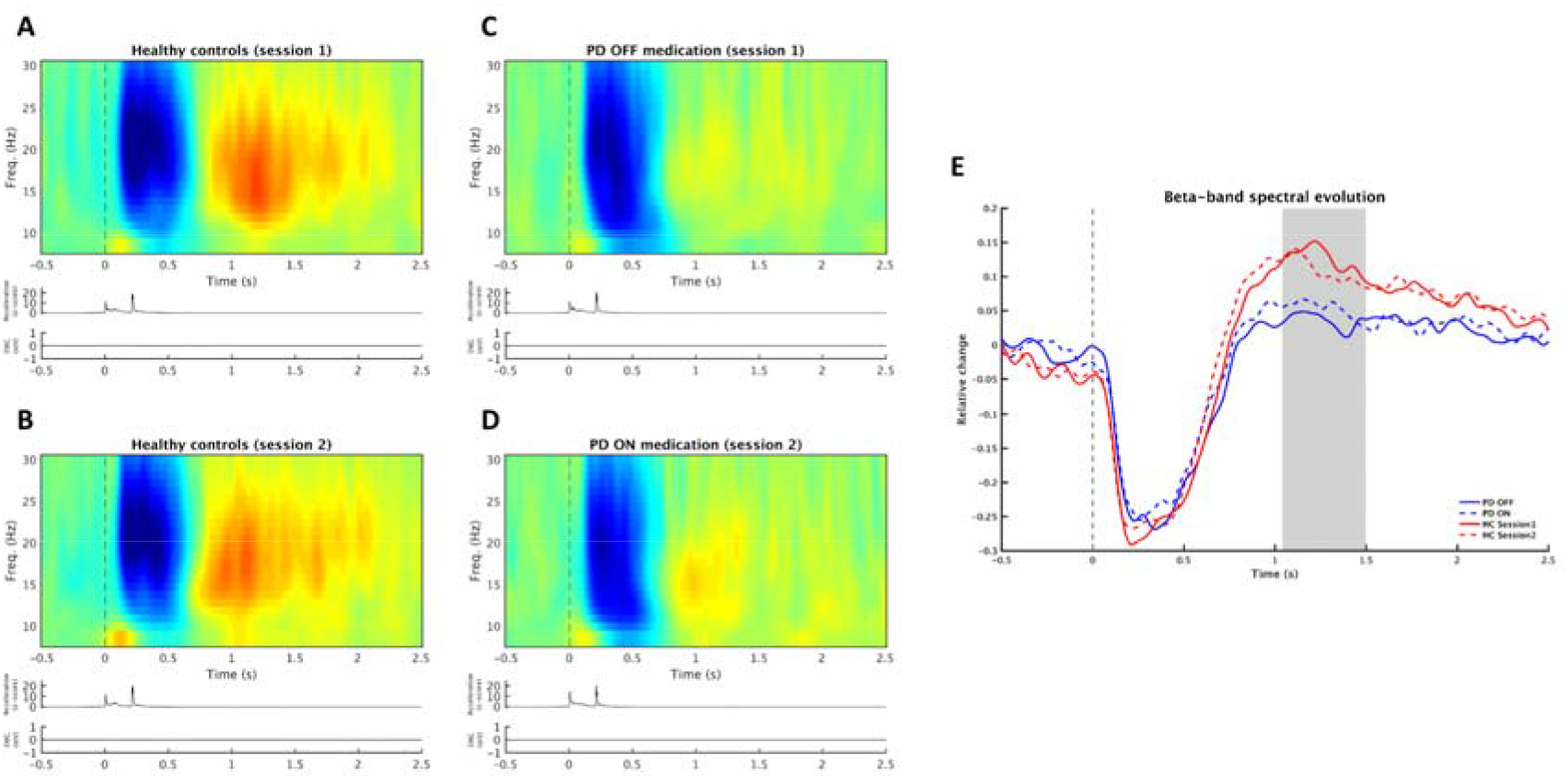
Beta-band response to proprioceptive stimulation. Time-frequency responses (TFR) of the cortical response to proprioceptive stimulation (movement onset at time=0) for HCs in session one (A) and session 2 (B), and PD patients OFF medication (C) and ON medication (D). Below each TFR are traces showing the acceleration of the movements and muscle activation measured with EMG. E) shows the temporal evolution of the beta-band for both sessions for both groups. The shaded area indicates the cluster that showed a significant difference between PD patients and HC.

The between-groups and between-sessions comparisons of the effect of levodopa on the beta/mu response to the proprioceptive stimulation yielded no significant result (p=0.45). Comparing the sessions for PD patients alone, without including the interaction between groups to correct for repetition effects, did not reveal any difference either (p=0.55).

### 3.4 Relation between beta rebound and baseline beta power

We compared the relationship between the average spectral power-change in the cluster covering the beta rebound described above and the average power spectral density from 1.2 s to 0.2 s before the onset of the passive movement, by comparing regression models with baseline power as a predictor (H_1_) and a model without baseline power (H_0_). The model comparison did not support the model containing baseline power over the model without it (BF_H1/H0_=0.83). Including interactions between baseline power, session, and group in the model (H2) and comparing it to the model without baseline power gave substantial evidence in favor of the model without baseline power (BF_H2/H0_=0.16).

## 4 Discussion

In the present study, we aimed at elucidating the processing of afferent proprioceptive information in PD by combining two different approaches. First, we used a computer-controlled proprioceptive stimulator that generates precisely controlled passive proprioceptive stimulation, with the aim of separating the processing of afferent proprioceptive information from that of efferent motor information. The controlled proprioceptive stimulation made sure the movements were identical for PD patients and HC in both sessions. Second, we studied PD patients ON and OFF Levodopa medication as compared to HC, with the aim of separating disease-related from medication-related effects. Our results show that when passive movements are used to generate proprioceptive stimulation, there is a definite difference in the cortical processing of afferent proprioceptive signals between PD patients and HC, manifested as a significant reduction—almost absence—of the beta rebound in PD patients as compared to HC (see Fig. 3). Our results also show that the beta rebound attenuation was not modulated by medication in PD, despite an evident effect from medication on overt motor symptoms, as assessed with MDS-UPDRS-III. The beta band attenuation hence emerges as a disease-related rather than medication-related change in the processing of afferent proprioceptive information in PD. Since medication does not modulate the attenuation, our results indicate that the disease-related change in proprioceptive processing does not directly reflect the dopaminergic networks of the brain.

The different stages in the cortical beta response to movements reflect different aspects in the processing of motor commands and proprioceptive feedback (Salmelin & Hari, 1994). The beta ERD, observed before and during movements (see Fig. 1), is taken to reflect a state of heightened sensitivity to efferent and afferent information within the motor system (Jenkinson & Brown, 2011). This notion has been supported by studies showing that reaction times to stimuli is negatively correlated with beta-band power, indicating that motor commands are executed more readily during an ERD when beta-band power is decreased (Heinrichs-Graham & Wilson, 2016; Shin et al., 2017). Such heightened sensitivity facilitates events such as motor commands being carried out efficiently as well as the integration of proprioceptive feedback while carrying out movements. The role of the beta rebound has been suggested to function an effective inhibition of motor responses (Pfurtscheller et al., 1996; Salmelin, Hämäläinen, Kajola, & Hari, 1995). As such the increase of beta oscillation during the beta rebound might reflect a “resetting” of the sensory-motor system, in terms of integrating the proprioceptive feedback from the action into the body schemata, thereby constructing an updated model of the position of the body and the limbs (Engel & Fries, 2010). The successful update of the body schemata is crucial for the calibration and execution of future actions as they will be dependent on the state of the body schemata to generate future motor commands and efferent signals.

The reduced beta rebound response in PD following proprioceptive stimulation might be understood as a deterioration in the processing and integration of proprioceptive singles: where errors in integrating proprioceptive signals lead to errors in the internal representation of body-state, and to more imprecise efferent motor commands. The reduction of the beta rebound in PD appears not just to be an after-effect of reduced beta ERS during active movements but related to the distinct processing of proprioceptive afferents.

Although the overt motor symptoms in PD patients changed upon Levodopa medication as compared to an initial OFF state (as rated by MDS-UPDRS-III), the attenuation of the beta rebound in PD patients did not change between Levodopa medication states. That we did not see any effect of medication on the beta band response to proprioceptive stimulation is in line with behavioral studies which showed that the error in detecting proprioceptive feedback for PD patients is not affected by Levodopa medication (Jacobs & Horak, 2006). Indeed, it has even been shown that Levodopa might worsen detection of proprioceptive feedback (Mongeon, Blanchet, & Messier, 2009; O’Suilleabhain, Bullard, & Dewey, 2001). Together with the behavioral findings, the finding that Levodopa medication did not alter the cortical responses to proprioceptive feedback results suggests that the dopaminergic system does not primarily mediate the later stages in the processing of proprioceptive signals. As an alternative, it has been proposed that processing of proprioceptive information in PD might rely less on the dopamine-dependent loop between basal ganglia, thalamus, and cortex, and instead involve pathways from the thalamus, through cerebellum to cortical areas (Wu & Hallett, 2013). Since MEG primarily detects activity from synchronous populations of pyramidal neurons in the cortex, we can, however, only speculate about the sub-cortical pathways responsible for propagating proprioceptive signals.

We acknowledge that it is possible that there might be effects of medication that we do not have sufficient statistical power to pick up due to our sample size. If there is a missed effect of Levodopa on the cortical processing of proprioceptive signals, these effects appear to be smaller than the difference in the beta-rebound response observed between PD patients and HC. Hence, even before we elucidate the post-movement beta rebound and its underlying mechanisms, this finding offers a potential marker for assessing the loss of proprioceptive function PD and disease progression in PD. The fact that the reduction of the beta rebound in the PD patients did not respond to levodopa medication suggest that it represents a reliable, disease-related measure that distinguishes PD patients from HC. Due to the relatively small sample size of PD patients in our current study, we cannot make a decisive conclusion about specific motor symptoms in PD. More studies with larger sample sizes are needed to better understand the role of the beta rebound—both in the general processing of sensory-motor signals and why it is attenuated in PD—and how the attenuation of the beta rebound is related to motor symptoms in PD.

Nevertheless, we can conclude is that at the cortical level, there appears to be a deficit in the processing of proprioceptive signals at the later cortical stage of processing and appears to be little affected by Levodopa medication.

## Acknowledgments

The experiment was carried out at NatMEG (http://natmeg.se; Karolinska Institutet, Stockholm, Sweden). The authors thank Robert Oostenveld for discussions and suggestions on the statistical analysis.

